# Novel non hot spot modification in Fks1 of *Candida auris* confers echinocandin resistance

**DOI:** 10.1101/2023.03.29.534846

**Authors:** Milena Kordalewska, Geselle Cancino-Prado, João Nobrega de Almeida Júnior, Igor Brasil Brandão, Renata Tigulini de Souza Peral, Arnaldo L. Colombo, David S. Perlin

**Author notes:** Address correspondence to Milena Kordalewska.

## Abstract

We determined echinocandin susceptibility and *FKS1* genotypes of thirteen clinical isolates of *Candida auris* recovered from four patients at a tertiary care center in Salvador, Brazil. Three isolates were categorized as echinocandin-resistant and harbored a novel *FKS1* mutation leading to an amino acid change W691L located downstream from hot-spot 1. When introduced to echinocandin-susceptible *C. auris* strains by CRISPR/Cas9, Fks1 W691L induced elevated MIC values to all echinocandins (ANF 16-32x; CAS >64x; MCF >64x).

---

Echinocandins are antifungal lipopeptides noncompetitively inhibiting β-1,3-D-glucan synthase that catalyzes synthesis of 1,3-β-glucan polymers of the fungal cell wall. In *Candida* spp., pharmacodynamic resistance to echinocandin class drugs occurs when a fungal strain develops mutations in two highly conserved hot-spot (HS) regions of the *FKS* genes encoding the catalytic subunits of β-1,3-D-glucan synthase. *In vitro*, presence of amino acid substitutions in Fks subunits manifests as elevated minimal inhibitory concentration (MIC) values (1).

*Candida auris* is an emerging fungal pathogen exhibiting elevated rates of antifungal drug resistance (including resistance to echinocandin class drugs) and causing hospital outbreaks (2). In *C. auris*, Fks1 HS regions are located at amino acid positions F635-P643 (HS1), and D1350-L1357 (HS2), and echinocandin resistance-conferring mutations were reported at several HS positions (**Table 1**). Here, thirteen isolates of *C. auris* (B61-B73), recovered from 4 patients (A, n= 4; B, n=7; C, n=1; X, n=1) at a tertiary care center in Salvador, Brazil, were evaluated for their echinocandin susceptibility and *FKS1* sequences. We performed antifungal susceptibility testing (AFST) with echinocandins (anidulafungin, ANF; caspofungin, CAS; micafungin, MCF) according to CLSI methodology (8, 9) with *C. parapsilosis* ATCC 22019 and *C. krusei* ATCC 6258 used as quality control strains. MICs were interpreted using tentative breakpoints as suggested by the CDC (10). Full *FKS1* gene was amplified by PCR, sequenced by Sanger sequencing, and its sequence analyzed (ref. gene B9J08_000964) as we described before (5). Three isolates recovered from urine of patient A were categorized as echinocandin-resistant (ANF=2-16 mg/l; CAS >16 mg/l; MCF >16 mg/l) and harbored a novel *FKS1* mutation G2072T leading to an amino acid change W691L (TGG→TTG). The remaining 10 isolates were echinocandin-susceptible (ANF MIC= 0.125-0.25 mg/l; CAS MIC = 0.125-0.25 mg/l; MCF MIC = 0.125-0.25 mg/l) and presented a WT *FKS1* genotype (**Table 2**). Full Fks1 sequences of clinical isolates B61-B73 were deposited at NCBI GenBank under accession numbers OQ632632-OQ632644.

**Table 1.**
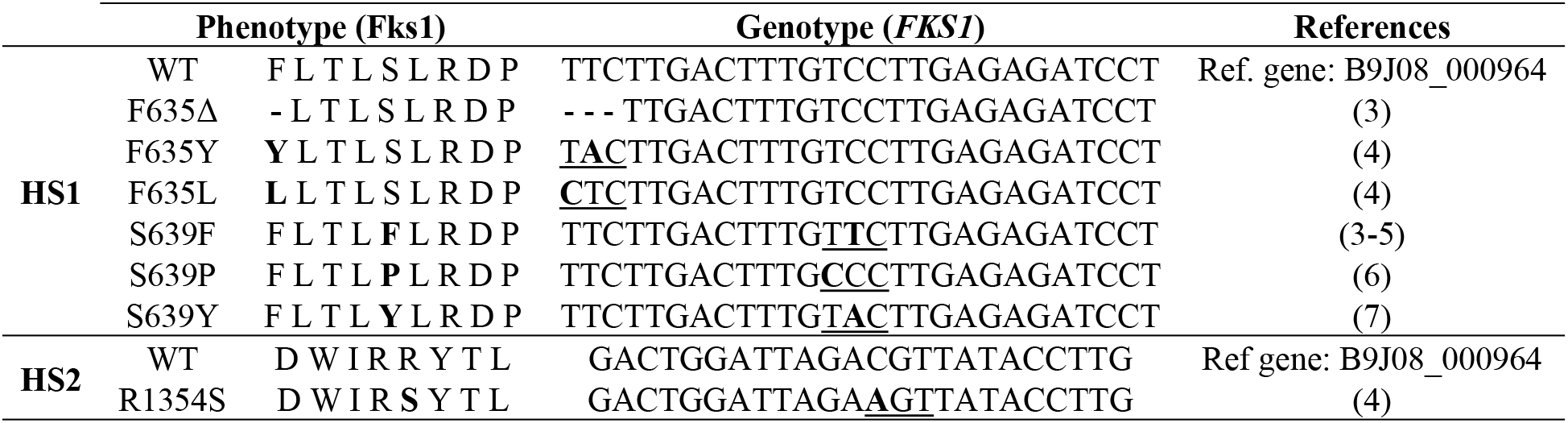
Echinocandin resistance-conferring alterations in Fks1 of *Candida auris*.

**Table 2.**
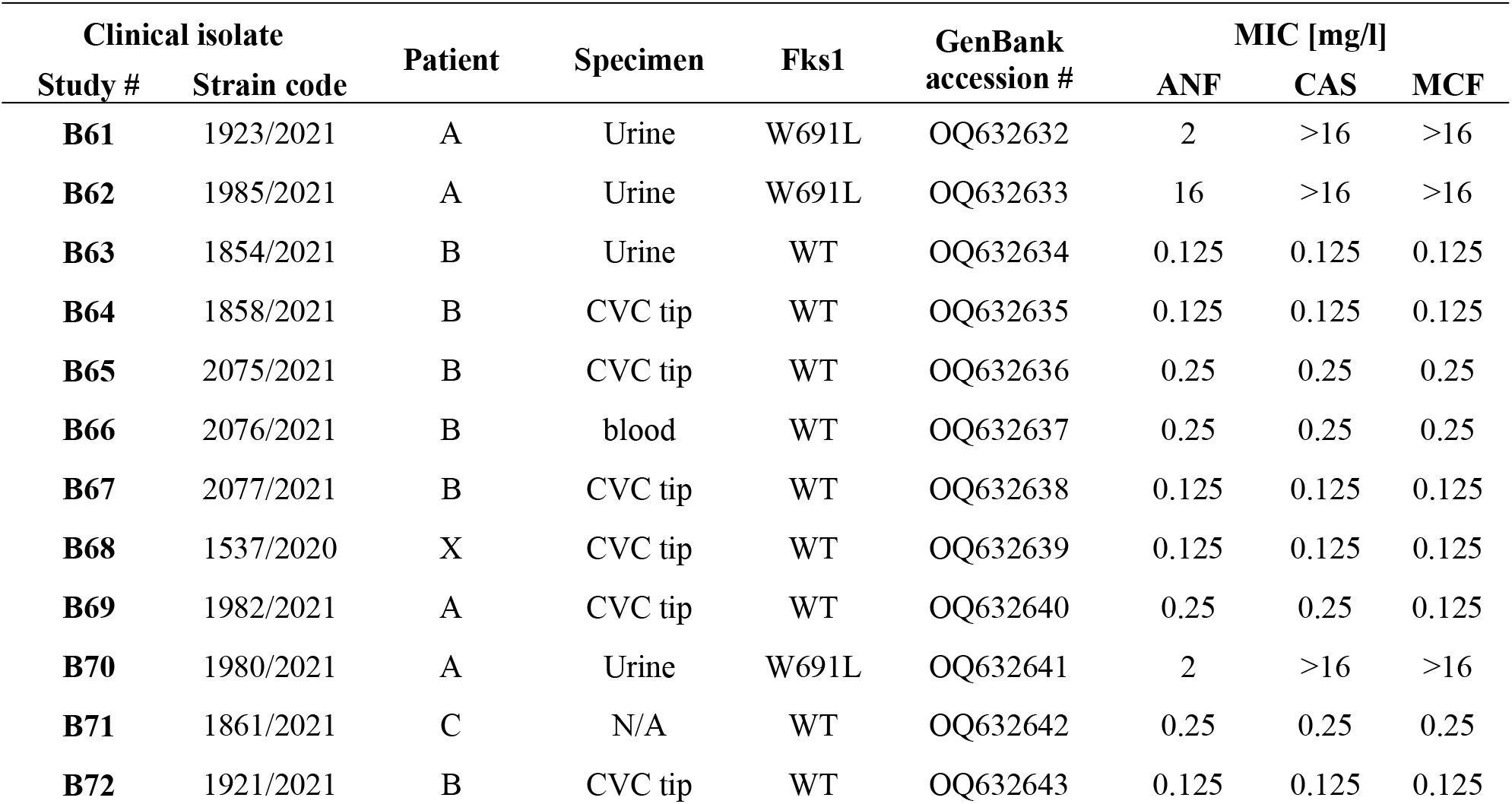
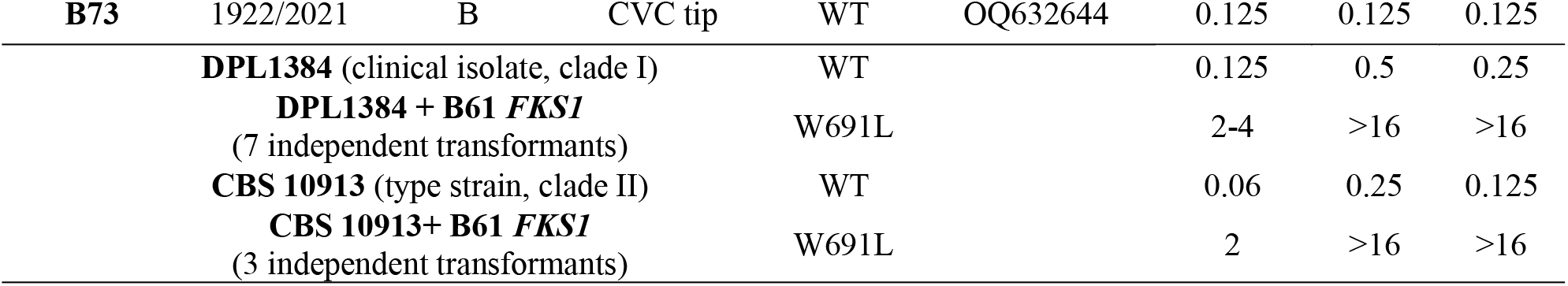
Antifungal drug susceptibility and Fks1 phenotypes of clinical isolates, parental strains and transformants.

W691 in Fks1 of *C. auris* is located 48 amino acids downstream from *FKS1* HS1 and is equivalent to W695 and W715 in *Saccharomyces cerevisiae’s* Fks1 and *C. glabrata’s* Fks2, respectively (11, 12). According to Johnson and Edlind, Fks1 W695 in *S. cereviasiae* is located in hot-spot 3 (AA 690 to 700) that is predicted to be embedded within the outer leaflet of the membrane and may interact with the extended aliphatic tails of echinocandins (11). In *C. glabrata*, W715L amino acid substitution in Fks2, was shown to lead to elevated echinocandin MIC values (12, 13).

To confirm that the Fks1 W691L variant confers echinocandin resistance, a wild-type (WT) *FKS1* gene fragment in echinocandin-susceptible *C. auris* DPL1384 (clinical isolate, clade I) and CBS 10913 (type strain, clade II) was replaced with a 465 bp *FKS1* fragment (C1899-C2364) harboring W691L (amplified from isolate B61) by using CRISPR/Cas9 system as described before (14), except that the cells were made electrocompetent with the Frozen-EZ yeast transformation kit (Zymo Research, Irvine, CA, USA). Repair template was PCR-generated with Q5 High-Fidelity DNA Polymerase (New England Biolabs, Ipswich, MA) and Cau_FKS1_CRISPR_F1 (5’-CTTCTTCTTGACTTTGTCCTTGAGAGATCCTATTAGAAACTTGTCCACC-3’) and Cau_FKS1_CRISPR_R1 (5’-GGTGCTCTCAAAGTTCTCTTGCCCTCGATTTCAGAAGGAACCTGGTGATAC-3’) primers. crRNA target sequence was CGATTTCAGAAGGAACCTGGTGG and Cau_FKS1_CRISPR_R1 primer introduced modified PAM site (without an amino acid change) into the PCR product to prevent Cas9 digestion of the repair template. The transformants were selected on YPD plates containing 4 mg/l MCF. Correctness of transformation was validated by PCR and sequencing of the entire *FKS1* gene (5). After that, the echinocandin MIC values of the correct transformants were determined according to CLSI methodology (8, 9). When introduced to an echinocandin-susceptible strains, *FKS1* W691L induced elevated MIC values to all echinocandins (ANF 16-32x; CAS >64x; MCF >64x) (**Table 2**). In conclusion, we report Fks1 W691L as a novel, echinocandin resistance-conferring non-hot-spot modification in *C. auris*. Our discovery emphasizes that despite significant progress in understanding echinocandin resistance in *C. auris* made in recent years, the diversity of *FKS1* mutations contributing to *C. auris* echinocandin resistance may be underestimated. Ultimately, improved knowledge of resistance mechanism will help diagnose and treat *C. auris*-infected patients.

## ACKNOWLEDGEMENTS

Partial results of this study were presented at ASM Microbe 2022 in Washington, DC. The study received Institutional Review Board approval CEP8260200319. J.A.N.J. and A.L.C. were awarded research grants (J.A.N.J.: FAPESP 2018/19347; A.L.C.: FAPESP 2017/02203-7) by Fundação de Amparo à Pesquisa do Estado de São Paulo. A.L.C. serves on advisory boards/educational programs of Amgen, Ache, Biotoscana-United Medical, Eurofarma, Mundipharma, Gilead, and Pfizer. D.S.P. receives funding from the U.S. National Institutes of Health (NIH) and contracts with Merck, Regeneron and Pfizer. He serves on advisory boards for N8 Medical and Scynexis. The remaining authors declare that the research was conducted in the absence of any commercial or financial relationships that could be construed as a potential conflict of interest.

